# Prostate Epithelial Genes Define Docetaxel-Sensitive Prostate Cancer Molecular Subtype

**DOI:** 10.1101/2020.08.26.269415

**Authors:** Hyunho Han, Kwibok Choi, Young Jun Moon, Ji Eun Heo, Won Sik Ham, Won Sik Jang, Koon Ho Rha, Nam Hoon Cho, Filippo G. Giancotti, Young-Deuk Choi

## Abstract

**BACKGROUND & OBJECTIVES:** Analysis of the transcriptomic landscape of prostate adenocarcinoma shows multidimensional gene expression variability. Understanding cancer transcriptome complexity can provide biological insight and therapeutic guidance. To avoid potential confounding factors, such as stromal contamination and stress-related material degradation, we utilized a set of genes expressed by prostate epithelial cells from single-cell transcriptome data of the human prostate gland.

**MATERIALS & METHODS:** Analyzing publicly available bulk and single-cell RNA sequencing data, we defined 1,629 genes expressed by prostate epithelial cells. Consensus clustering and CIBERSORT deconvolution were used for class discovery and proportion estimate analysis. The Cancer Genome Atlas Prostate Adenocarcinoma (TCGA-PRAD) dataset served as a training set. The resulting clusters were analyzed in association with clinical, pathologic, and genomic characteristics and impact on survival.

**RESULTS:** TCGA-PRAD tumors were separated into four subtypes: A (30.0%), B (26.0%), C (14.7%), D (4.2%), and mixed (25.0%). Subtype A was characterized by low frequency of ETS-family fusions and high expression of *KLK3*, which encodes prostate-specific antigen (PSA). Subtype B showed the highest expression of *ACP3*, encoding PAP (prostatic acid phosphatase). Subtypes C and D were commonly associated with advanced T/N stages, high Gleason grades, and p53 or PIK3CA mutations. In silico drug-sensitivity screening suggested that subtype B is likely sensitive to docetaxel and paclitaxel. Serum PSA/PAP ratio was predictive of a radiographic response to docetaxel in metastatic castration-resistant prostate cancer patients.

**CONCLUSION:** We propose four prostate adenocarcinoma subtypes with distinct transcriptomic, genomic, and pathologic characteristics. PSA/PAP ratio in advanced cancer may aid in determining which patients would benefit from maximized androgen receptor inhibition or early use of antimicrotubule agents. Molecular subtypes and biomarkers must be validated in a prospective cohort study.

## INTRODUCTION

Stages and grades of cancers are linear-scale indicators to predict patient survival and determine treatment options. For prostate cancer (PCa), TNM staging, Gleason score, and serum prostate-specific antigen (PSA) level have been extensively utilized. Recently, molecular genetics provided another level of resources (i.e., genomic scoring systems based on the expression of multiple genes) [1,2]. However, evidence suggests that cancer variability is multidimensional. Transcriptomic analysis of breast cancer revealed molecular subtypes of tumors—luminal A/B, ERBB2+, and basal-like [3,4]. This revolutionized clinical practice and basic research in breast cancer, which has been translated to other cancer types [5]. In urologic tumors, transcriptomic subtyping identified that urothelial carcinomas likely respond to immune checkpoint inhibitors [6].

There have been several attempts to subtype PCa, including ETS transcription-factor–based classifications and luminal/basal lineage models directly adapted from breast cancer [7–9]. However, the utility of these approaches was limited because they provided no clinical information beyond known risk factors. Further, technical issues remain to be resolved. Whole-transcriptome analysis is susceptible to potential confounding factors, such as stromal contamination, and PCa is characterized by multifocality and/or intratumoral heterogeneity [10,11]; therefore, it is likely that a tumor may be composed of more than two molecular subtypes that differ in the tumor cell, as well as tumor-microenvironment gene expression [12–14].

Understanding of the tumor transcriptome has recently been enhanced by single-cell RNA-sequencing, which revealed that molecular subtypes can be further dissected [15], and that heterogeneity exists as a spectrum of potentially interchanging states [16]. For prostate tissue, single-cell analysis precisely defined epithelial-expressed genes and confirmed the existence of luminal, basal, or bipotential progenitor populations with specific anatomical locations and potential relevance to cancer characteristics [17–19].

Thus, we hypothesized that the prostate adenocarcinoma transcriptome can be interpreted based on the cell-of-origin of gene expression. Using prostate epithelial-cell–expressed genes from single-cell and mass transcriptome data, we developed a single-sample subtype classifier with proportion estimate (PE) for tumor RNA-Seq data based on an established deconvolution analysis tool. We report four transcriptomic subtypes with potential therapeutic relevance to antimicrotubule agents and inhibition of androgen signaling, as well as utility of serum biomarkers PSA and prostate-specific acid phosphatase (PAP) to identify the subtypes.

## MATERIALS AND METHODS

### Prostate Epithelial-Expressed Gene Identification from Single-Cell and Bulk RNA-Seq Data

We used *single-cell RNA-Seq data* from Henry et al [17] that used three human prostate specimens. Mapped read count data were downloaded from GEO (GSE117403) and aggregated using 10X Genomics Cell Ranger aggregate function. We followed the analysis pipeline of Henry et al and replicated differentially expressed gene (DEG) lists for luminal, basal, club-like, hillock-like, and neuroendocrine prostate epithelial cells. We analyzed the *bulk RNA-Seq data* of corresponding fluorescence-activated cell sorting-isolated human prostate cell types from the same study [17]. We downloaded the fragments per kilobase million (FPKM)-value matrix of basal, luminal, and other epithelia, and fibromuscular stroma from GEO (GSE117271), selecting genes overexpressed by more than 5-fold by all epithelia, or by an epithelial subpopulation vs. the remainder of the epithelia. We merged the DEG lists from single-cell and bulk datasets, and kept genes that mapped to an Entrez gene ID.

### Consensus Clustering of TCGA-PRAD (The Cancer Genome Atlas Prostate Adenocarcinoma) RNA-Seq Data

Consensus clustering is a resampling-based method for class discovery and visualization of gene expression array data [20]. We downloaded annotated TCGA-PRAD gene expression, clinical, and genomic data from the UCSC Xena browser. RNA-Seq by expectation maximization (RSEM) data containing mRNA expression levels of 550 samples were uploaded to the GenePattern Public server (cloud.genepattern.org). We used Consensus Clustering Module version 7.2 with parameters set at: K_max_=15; resampling iterations=20; clustering algorithm=self-organizing map; cluster by=columns; distance measure=Euclidean; resample=subsample with a proportion of 0.80; merge type=average; descent iterations=2,000; normalize type=row-wise; normalization iterations=0. We used pre-calculated DNA purity (ABSOLUTE, CLONET) and RNA purity (ISOpure, DeMix purity) scores, AR activity score, and AR mRNA and protein expression from the TCGA-PRAD dataset at the cBioPortal [21]. Survival data from the TCGA Pan-Cancer Atlas were also downloaded from the cBioPortal.

### RNA-Seq Data Deconvolution and Single-Sample PE

We used CIBERSORT, a digital cytometry tool for deconvolution of heterogeneous tissues based on bulk mRNA-Seq data [22]. It is based on a linear support vector-regression model of a sum of pure tissue- or cell-type–specific reads of all cell types, weighted by the respective cell-type proportions in a given dataset. RNA-Seq read-normalized gene expression values (RSEM, RPKM, and FPKM for TCGA, CPC-GENE and DKFZ, and SU2C-PCF datasets, respectively) with Entrez gene ID and HUGO gene-symbol annotations were loaded as a “mixture” file. The gene signature was defined by the average gene expression values of prostate epithelial-expressed genes in the TCGA-PRAD dataset clusters predetermined by Consensus Clustering.

### In silico Docetaxel and Paclitaxel Sensitivity Test

To predict docetaxel sensitivity in PCa patients, we used a previously published gene expression signature associated with breast tumor response to docetaxel therapy, evaluated by the degree of reduction in tumor size [23]. Genes overexpressed (n=13) and underexpressed (n=43) in docetaxel responders were used as gene sets to run single-sample gene-set enrichment analysis (ssGSEA) loaded as a module in the GenePattern platform. The docetaxel responder score was calculated by subtracting the ssGSEA score of underexpressed genes from that of overexpressed genes. To predict paclitaxel sensitivity, we searched the cancer therapeutics response portal (CTRP v2, http://portal.broadinstitute.org/ctrp.v2.1) and selected genes whose expression correlated positively (Pearson r>0.3) with cancer cell lines relative sensitivity to paclitaxel (1-area under curve value). All score values underwent z-score normalization.

### Docetaxel Response Analysis in the Yonsei University Health System (YUHS) Database

The study design was approved by Severance Hospital Institutional Review Board (IRB #4-2020-0652). The YUHS Big-Data team identified cases of metastatic castration-resistant prostate cancer (mCRPC) patients who: 1) underwent at least three consecutive cycles of docetaxel-predisone chemotherapy, 2) had abdominopelvic CT and whole-body bone scan imaging before, during, and after chemotherapy to assess radiographic response, and 3) had serum PSA and PAP test results acquired within 30 days ahead of initial chemotherapy start date. Docetaxel response was measured using RECIST 1.1 criteria.

### Statistics and reproducibility

Statistical analyses were performed with GraphPad Prism version 8.4.3 (GraphPad, San Diego, CA, USA). P-values were estimated using log-rank (Mantel-Cox) test for survival curve comparison, unless indicated otherwise. For analysis of correlation between drug-sensitivity scores and subtype PEs, Spearman r values and two-tailed P-value were reported. For multiple comparisons, ANOVA and Kruskal-Wallis test were used.

## RESULTS

### Prostate Epithelial-Cell–Expressed Genes Define Four Tumor Clusters

We extracted lists of DEGs from single-cell and bulk RNA-Seq data of human prostate-tissue samples, identifying 1,629 genes expressed by epithelial cell populations vs. all other cell types in the sample (Table S1, courtesy of Dr. Douglas Strand at UT Southwestern Medical Center) [17]. Initial consensus self-organizing map clustering of the TCGA-PRAD dataset using 1,593 genes (24 removed due to gene ID/symbol mismatching) suggested an optimal number of clusters in the range 2–5 (Figure S1), but the absence of a plateau in consensus cumulative distribution function (CDF) plots implied that the population could not be cleanly separated. Alternatively, we filtered samples by DNA purity (ABSOLUTE, CLONET purity values >0.5) and RNA purity (ISOpure, DeMix purity values >0.5), selecting 138 of 275 samples of purity data available (50.2%, Table S2). Using this “pure” subset, we repeated consensus clustering and found four robust clusters (clusters A–D) with a minimal proportion of ambiguously clustered pairs (Figure 1A-D).

**Figure 1.**
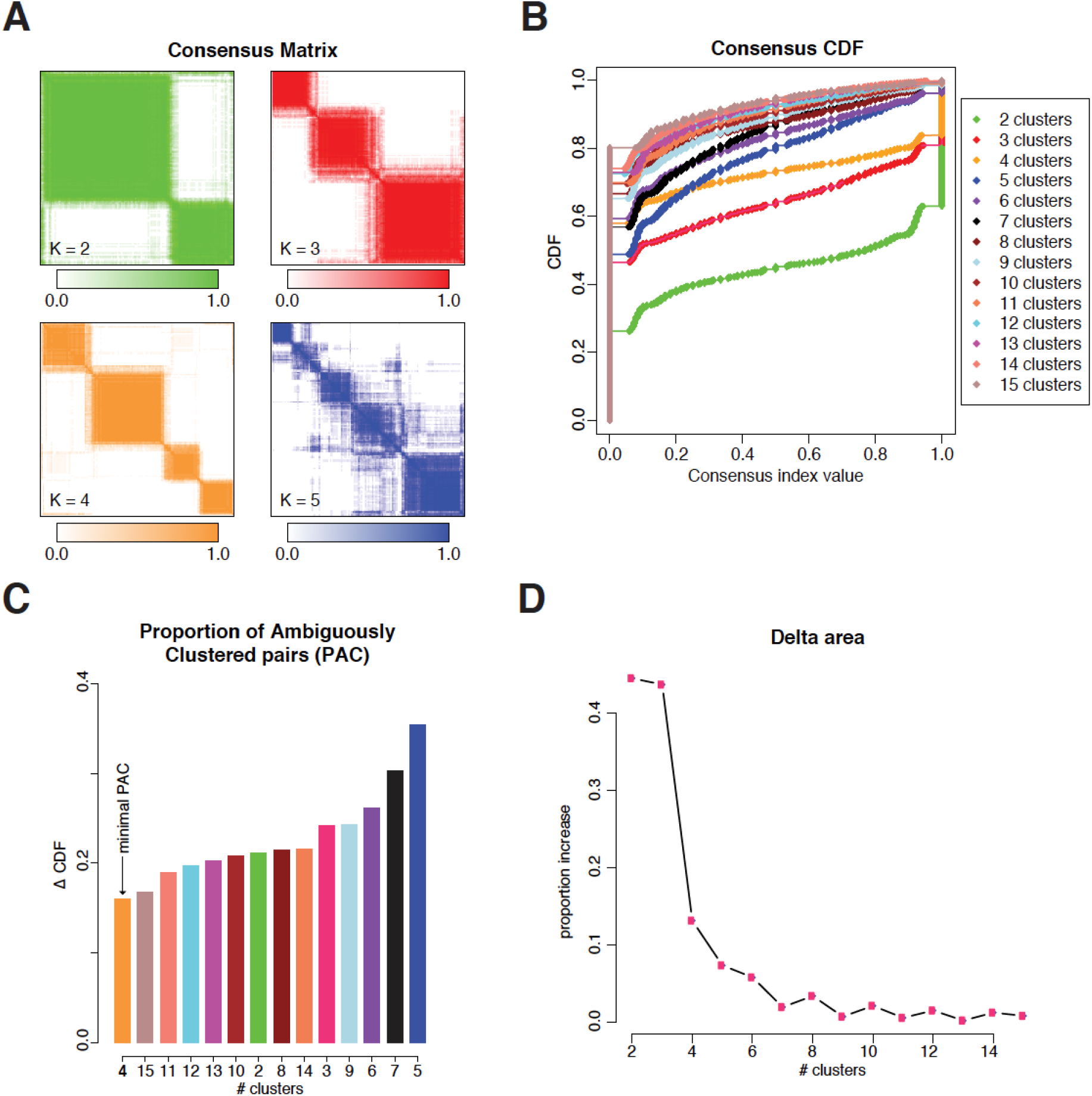
Consensus Clustering of The Cancer Genome Atlas Prostate Adenocarcinoma (TCGA-PRAD) RNA-Seq Data Filtered by DNA and RNA Purity. **A.** Heatmaps of clustered consensus matrices of K=2 to K=5. **B.** Consensus cumulative distribution function (CDF) plots of K=2 to K=15. **C.** Proportion of ambiguously clustered pairs (PAC), represented by Δ CDF_K_ (CDF_K_[index value 0.9]-CDF_K_ [index value 0.1]). Arrow indicates minimal PAC at K=4. **D.** Delta area plot showing the proportion increase in area under the CDF curve comparing K and K-1.

### Genomic Characteristics of Clusters Assigned by Deconvolution Analysis

We generated gene signatures containing 1,271 DEGs among the four clusters and calculated PEs of each cluster in the TCGA dataset by running CIBERSORT deconvolution analysis (P<0.05) (Figure 2A, Table S3). A sample was assigned to each cluster when the cluster PE was >0.5, and those with maximal PE≤0.5 were designated “mixed”. By this definition, samples were classified as cluster: A (n=163, 30.0%), B (n=141, 26.0%), C (n=80, 14.7%), D (n=23, 4.2%), or mixed (n=136, 25.0%). We analyzed the genomic characteristics of clusters A–D using the cBioPortal’s group comparison function [24]. Cluster A was characterized by frequent SPOP mutation and chr6q21 homodeletion, and the absence of ETS-family fusion. Cluster B was characterized by PTEN deletion and ETS fusion. In addition, clusters C and D showed further alterations, including TP53 mutation/loss of heterozygosity, PIK3CA mutation, and amplifications of chr8q24.3, which harbors FAK (*PTK2*, encoding focal adhesion kinase) (Figures 2B, C and S2A). Compared to previously identified TCGA-PRAD molecular subtypes, cluster A consisted of alterations in SPOP and other genes; cluster B, alterations purely in ERG; cluster C, in ERG and ETV1; and cluster D, in ERG and ETV4 (Figure 2D).

**Figure 2.**
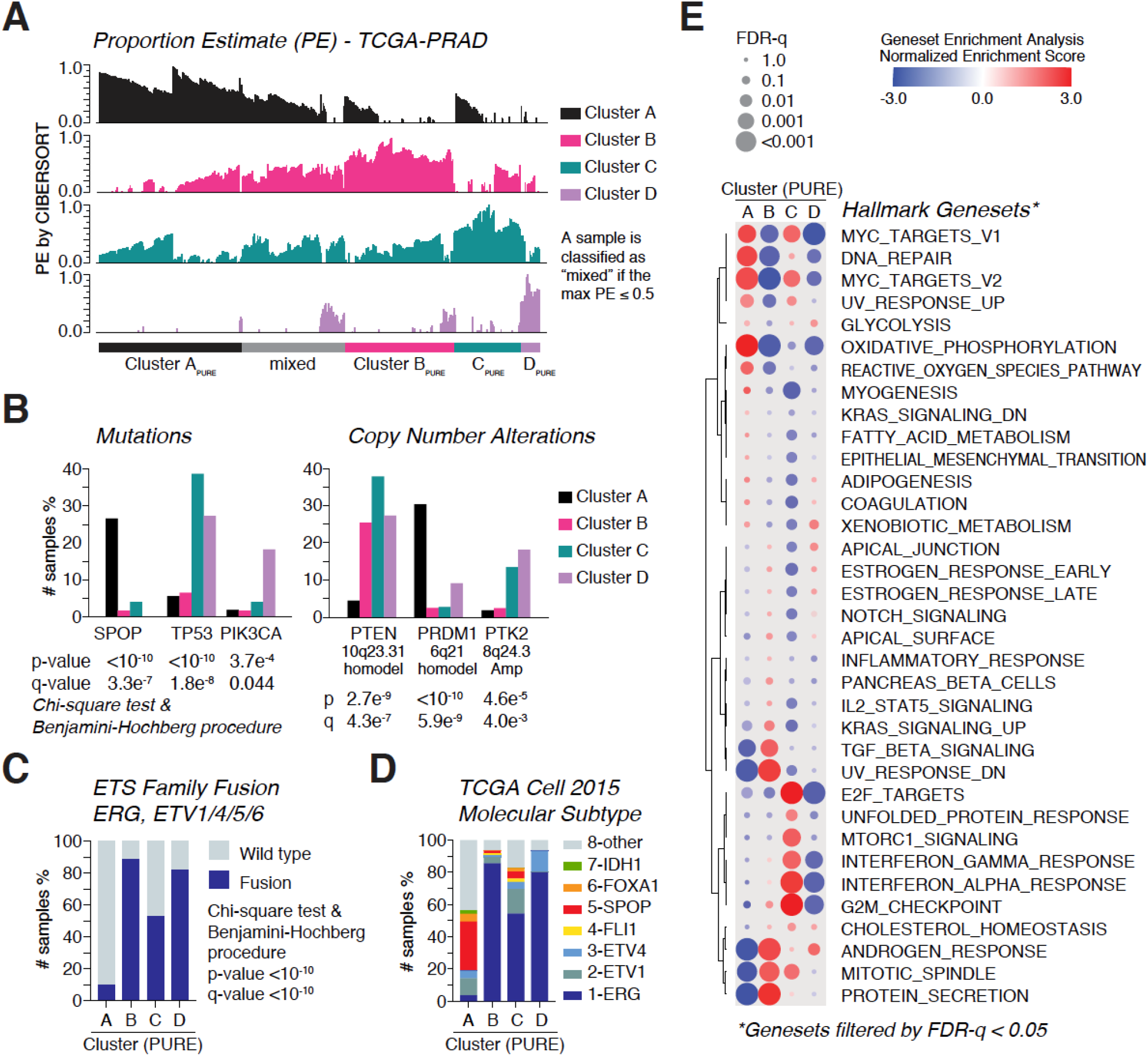
Genomic and Transcriptomic Characteristics of the Four Clusters from the TCGA-PRAD Data. **A.** Proportion estimate (PE) bar graphs of the four clusters from the CIBERSORT deconvolution analysis of the TCGA-PRAD dataset. A sample was classified as “PURE” if PE_max_ >0.5. The sample was classified as “mixed” if PE_max_ ≤0.5. **B.** Cluster-wise frequencies of SPOP, TP53, and PIK3CA mutations and copy-number alteration events (i.e., chromosome 10q23.31 homodeletion (represented by PTEN), chromosome 6q21 homodeletion (represented by PRDM1), and chromosome 8q24.3 amplification (represented by PTK2)). P-values and Q-values were calculated by chi-square test and Benjamini-Hochberg procedure. **C.** Cluster-wise frequencies of ETS-family fusions (ERG; ETV1, 4, 5, or 6). **D.** Cluster-wise distribution of the eight molecular subtypes reported by the TCGA in 2015. **E.** Dot plot of enrichment results for each cluster vs. the remaining clusters for comparison. Results of analysis of the hallmark 50 gene sets of minimal FDR-Q <0.05 are shown.

### Transcriptomic Characteristics and Cluster Androgen Receptor (AR) Activity

We performed GSEA comparing each cluster with the other three using the hallmark 50-gene sets [25] (Figure 2E). Cluster A was enriched in MYC_TARGETS, DNA_REPAIR, and OXIDATIVE_PHOSPHORYLATION. Cluster B was enriched in TGF_BETA_SIGNALING, ANDROGEN_RESPONSE, and PROTEIN_SECRETION. Cluster C was enriched in E2F_TARGETS, MTORC1_SIGNALING, and INTERFERON responses. Cluster D was enriched in GLYCOLYSIS and ESTROGEN responses (Figure 2E). AR activity score was highest in cluster A, but *AR* mRNA levels were low in cluster A and D. AR protein levels were not significantly different among the clusters (Figure S2B-D).

### Pathologic and Serum-Marker Characteristics of Clusters

Gleason score and pathologic T and N stage distributions were significantly different among clusters (Figure 3A-C). Clusters C and D were characterized by a higher Gleason score and more advanced T and N stages than A and B. In contrast, preoperative (radical prostatectomy, RP) serum PSA levels were not significantly different among clusters, either by average level or distribution (Figure 3D). We further examined PSA levels of clusters A and B. When stratified by pT stage, pT2C cluster A tumors showed significantly higher PSA levels than pT2C cluster B. Indeed, expression of *KLK3* mRNA, which encodes PSA, was significantly higher in cluster A than in B or the other two clusters (Figure 3F). In contrast, the level of ACP3 mRNA, encoding prostatic acid phosphatase (PAP), was highest in cluster B (Figure 3G). Consequently, the ratio of *KLK3*/ACP3 gene expression was lowest in cluster B (Figure 3H). Serum PAP level data was not available for TCGA samples.

**Figure 3.**
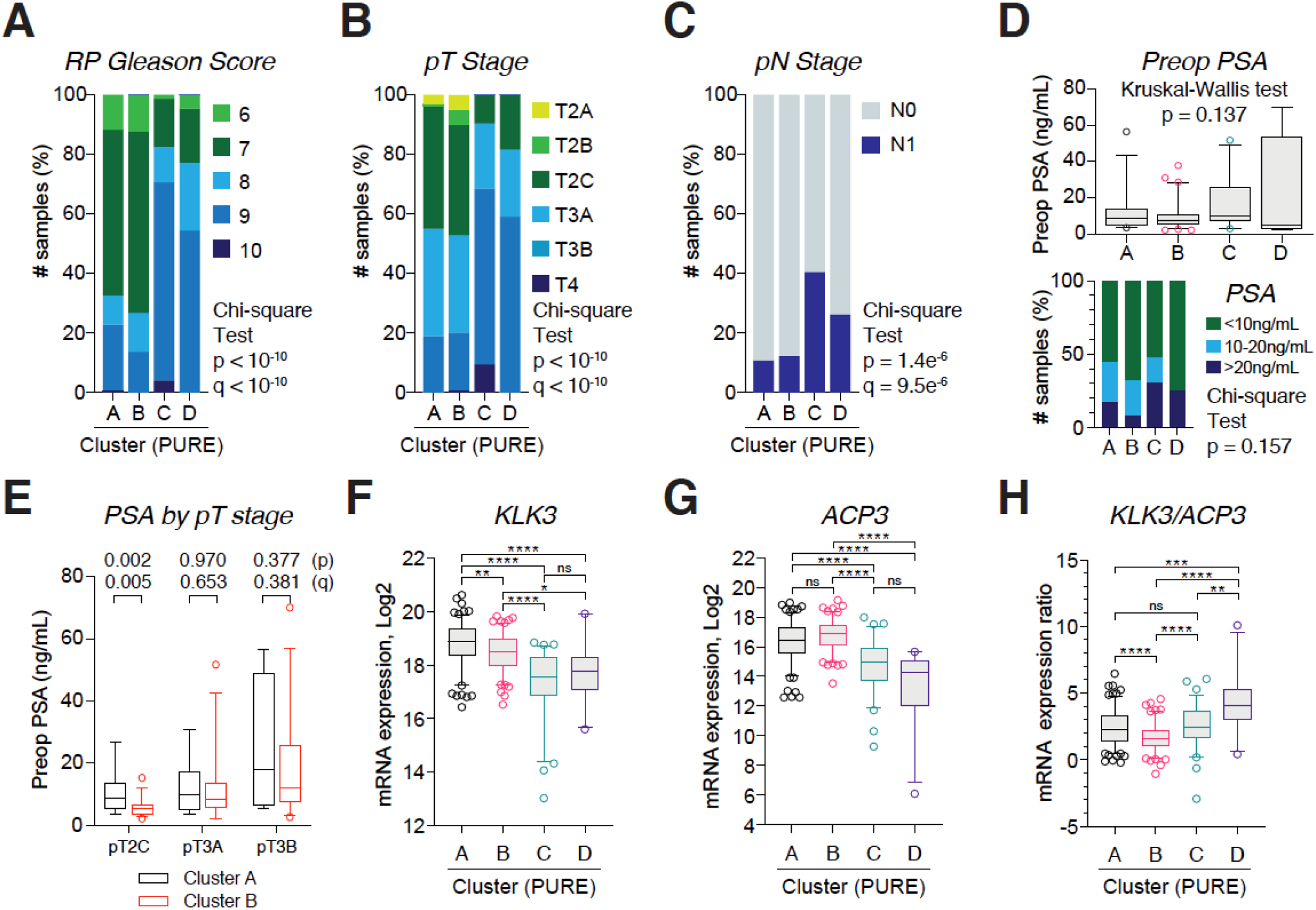
Pathologic and Biomarker Characteristics of the Four Clusters. **A-C.** Cluster-wise distributions of radical prostatectomy (RP) Gleason score (A), pathologic T (pT) stage (B) and pN stage (C). P-values and Q-values were determined by chi-square test and Benjamini-Hochberg procedure. **D.** Preoperative (RP) serum prostate-specific antigen (PSA) levels presented in box plot (5%–95%) and stacked bar chart (class interval: 10 ng/mL, 20 ng/mL). **E.** Cluster A and B preoperative serum PSA levels stratified by pT stage. Multiple *t*-test, Holm-Sidak method, without assuming a consistent standard deviation. **F-H.** *KLK3*, ACCP and *KLK*/ACP3 ratio mRNA expression (log2) levels among the clusters. RNA-Seq by expectation maximization (RSEM; batch-normalized from Illumina HiSeq_RNASeqV2) (log2) values were downloaded from the cBioPortal. Multiple comparison assessed by Kruskal-Wallis test.

### Prognostic Value of Clusters

Despite variations in pT/N stages and Gleason scores, cluster survival curves were not well separated, either by disease-free survival or progression-free survival (Figure 4A, B). Only the cluster B/C comparison reached statistical significance. When impact of known prognostic variables was assessed in each cluster, primary Gleason grade (grade 3 vs. 4) showed prognostic significance in clusters A, B, and D, but not C (Figure 4C).

**Figure 4.**
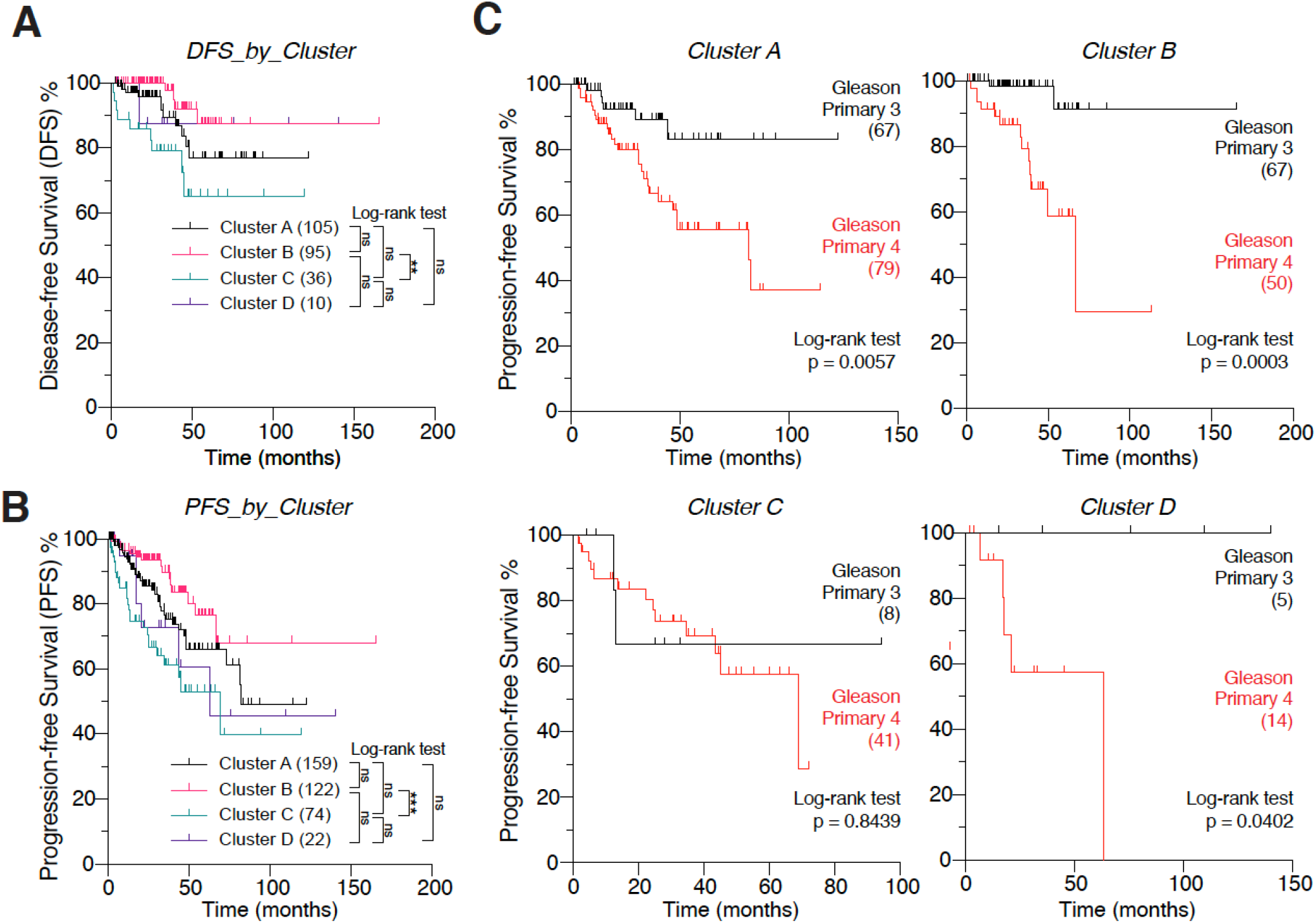
Prognostic Value of the Four Clusters. **A** and **B.** Disease-free survival (A) and progression-free survival (B and C) curves of the four clusters. P-values by log-rank test. Multiple comparison by Bonferroni correction. ns=not significant. *p<0.05; **p<0.01; ***p<0.001. (n of samples). Survival data from the TCGA-PRAD Pan-Cancer Atlas. **C.** Progression-free survival of RP primary Gleason grade 3 vs. grade 4, stratified by cluster. P-value by log- rank test.

### Application in Additional Prostate-Cancer Datasets

We applied the gene signature for deconvolution of three additional PCa datasets. The CPC-GENE 2017 dataset consists of localized non-indolent tumors (Gleason score 6–7, clinically organ-confined) [26]. The DKFZ 2018 dataset consists of tumors diagnosed in patients <50 years old [27], and the SU2C-PCF 2019 dataset consists of mCRPCs [28]. In the three datasets, the distribution of Gleason score, pT stage, ETS fusion, and SPOP and TP53 mutation events were similar to that of the TCGA dataset, confirming cluster genomic and pathologic characteristics (Figure 5A-C). Of note, clusters B and A were the largest groups in the CPC-GENE and DKFZ datasets, respectively. In the mCRPC dataset, tumors were assigned to only cluster A or C or designated “mixed” (Figure 5C). Serum PSA level was available in the SU2C_PCF dataset, and cluster A tumor PSA level was higher than that of cluster C tumors (P<0.01). This was after excluding samples with histologic neuroendocrine features, which are associated with extremely low serum PSA levels [29].

**Figure 5.**
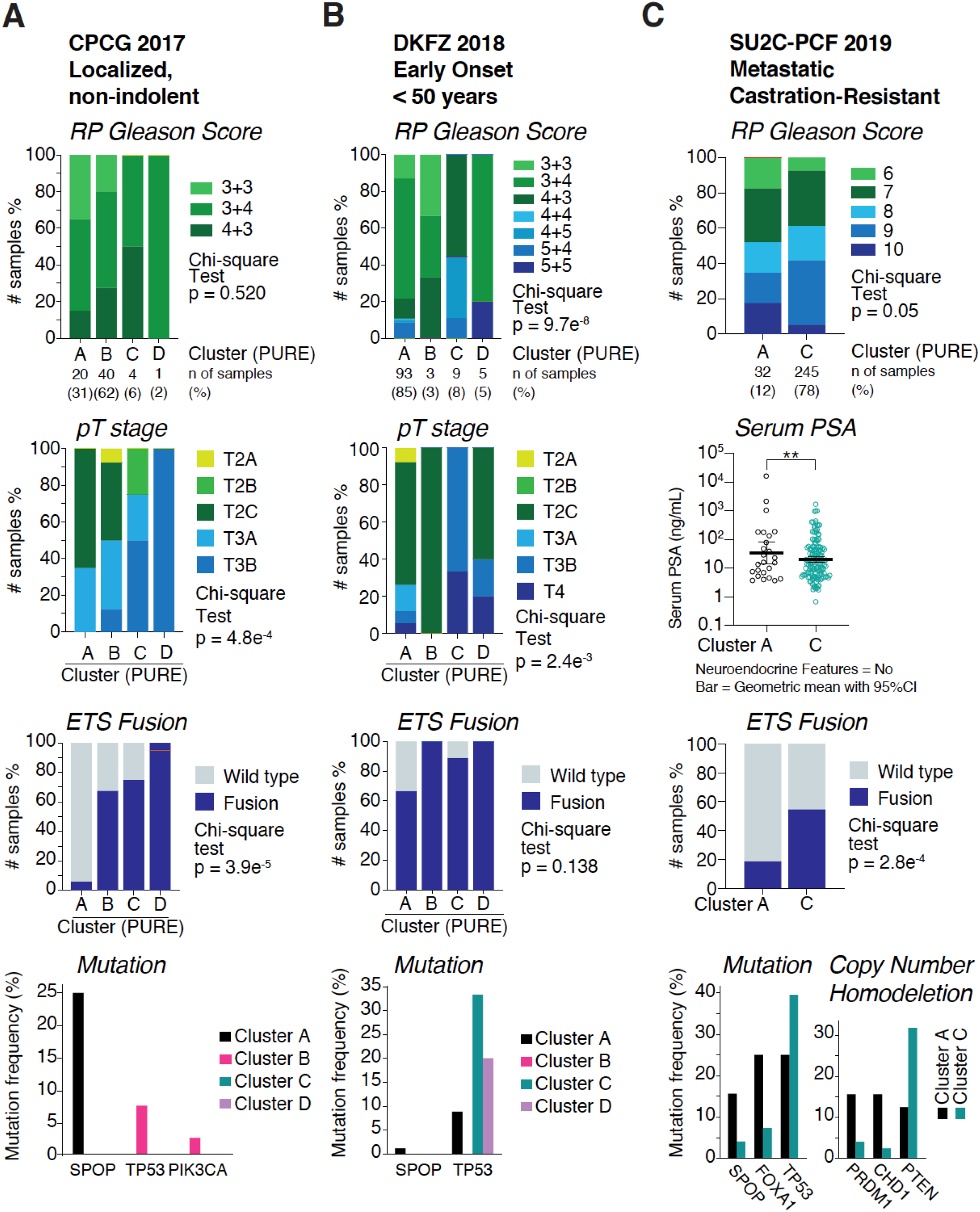
Application of Single-Sample Classifier in Additional Prostate Cancer Datasets. **A-C.** RP Gleason score, pT stage distributions, and frequencies of ETS-family fusions and gene mutations among clusters identified in three additional prostate cancer datasets. The CPC-GENE 2017 dataset consists of localized non-indolent tumors (Gleason score 6–7, clinically organ-confined). The DKFZ 2018 dataset consists of tumors diagnosed in patients <50 years of age. The SU2C-PCF 2019 dataset consists of metastatic castration-resistant tumors (mCRPCs). For SU2C-PCF dataset, samples of concordant deconvolution analysis results between RNA-Seq Poly-A and Capture were included. The dataset provided serum PSA (measured at mCRPC state) and copy-number alteration data.

### In silico Docetaxel and Paclitaxel Sensitivity Test

In silico drug-screening can identify drugs potentially effective in a cancer model based on its transcriptome data [30]. We used a modified version of the in silico drug-screening logic to predict sensitivity to docetaxel and paclitaxel in PCa samples. Briefly, gene sets were defined based on genes reported to positively or negatively correlate with tumor drug response. Drug response or sensitivity score in a sample is then computed by ssGSEA. Using this algorithm, we found that cluster B is the most likely to be a docetaxel responder (Figure 6A, Table S4). Correlation analysis was consistent with cluster-wise comparison, showing that cluster B PE positively correlates, and cluster A PE negatively correlates, with docetaxel responder score (Figure 6B). The analysis also predicted that cluster A is least likely to be sensitive to paclitaxel (Figure 6C), and cluster A PE negatively correlated with the paclitaxel sensitivity score (Figure 6D).

**Figure 6.**
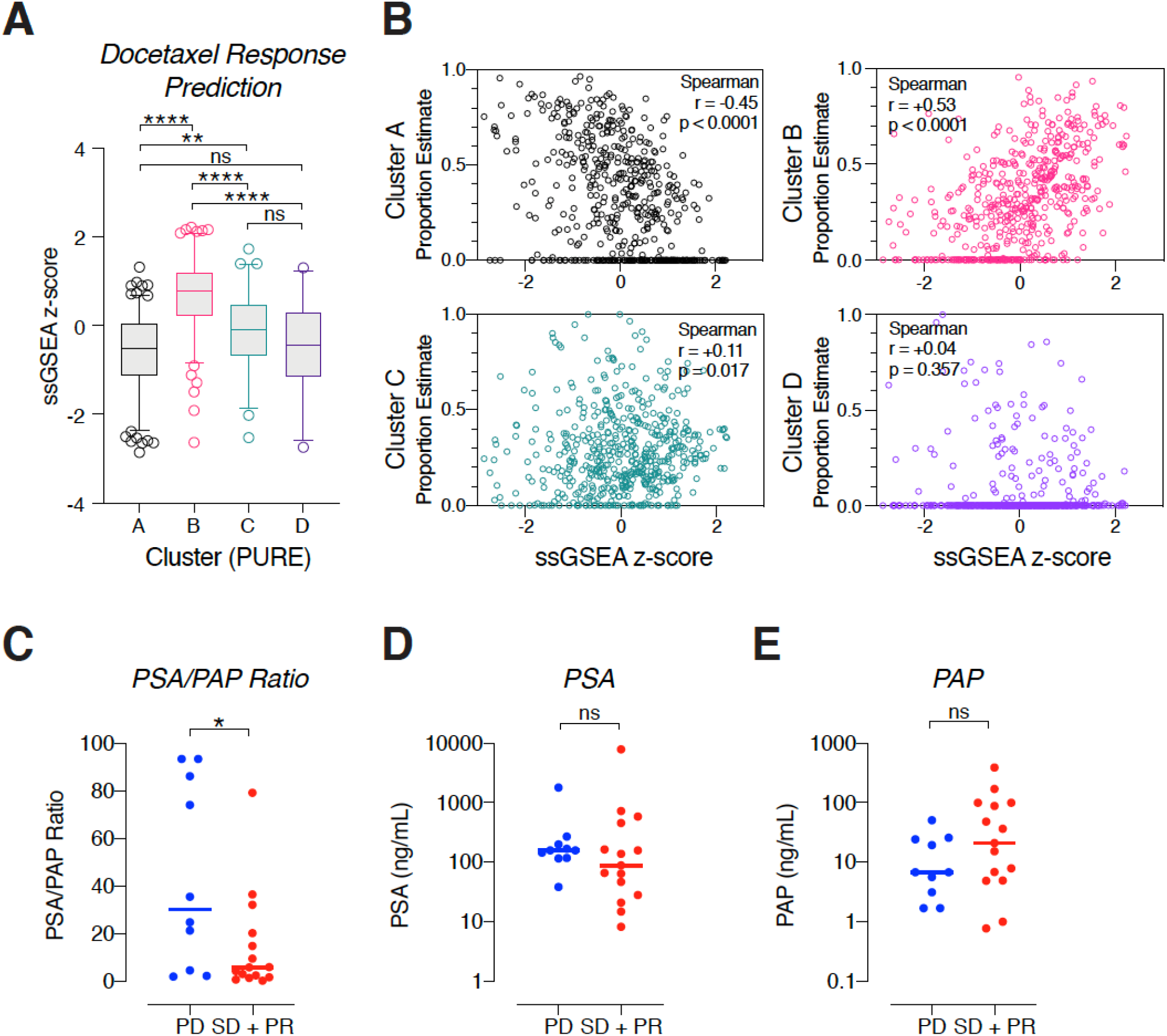
In silico Docetaxel Sensitivity Test and Validation in Patients. **A.** Pair-wise comparison of docetaxel response score among the four clusters. Kruskal-Wallis test and multiple comparison by Dunn’s multiple comparisons test. ns=not significant. *p<0.05; **p<0.01; ***p<0.001; ****p<0.0001. **B.** Scatter plots of the four cluster PEs (Y-axis) and docetaxel response score of each sample. Spearman correlation coefficient and p-values are shown in the box. **C-E**. Pre-treatment serum PSA/PAP ratio and absolute levels in docetaxel responder and nonresponder groups treated at Yonsei University College of Medicine between 2005–2020. PD=progressing disease; SD=stable disease; PR=partial response by RESIST 1.1 criteria. ns=not significant. *p<0.05.

### PSA/PAP Ratio in mCRPC is Associated with Docetaxel Response

Based on the *KLK3*/ACP3 gene expression ratio and in silico drug sensitivity results, we hypothesized that serum PSA/PAP ratio can predict the docetaxel response of mCRPC patients. We assumed that: 1) serum PSA/PAP ratio is reflective of tumor cell *KLK3*/ACP3 gene expression, and 2) serum PSA/PAP ratio remains constant despite the effect of LHRH agonists and AR blockers on absolute levels. We found 25 cases eligible for response analysis. Mean pre-treatment PSA and PAP levels were 546.6±1549.6 ng/mL (min. 8.2; max. 7929) and 45.7±81.4 ng/mL (min. 0.77; max. 390), respectively. Ten patients were non-responders (immediate progressive disease) and fifteen patients were responders (stable disease, 13 cases; partial response, 2 cases/responses seen within at least 3 months during/after treatment). Mean responder PSA/PAP ratio was significantly lower than that of non-responders (14.5 vs. 43.8, P<0.05, unpaired *t*-test with Welch’s correction) (Figure 6C). Mean PSA and PAP levels were not significantly different between responders and non-responders (Figure 6D, E).

## DISCUSSION

The pioneering research of Perou et al. provided the groundwork for transcriptome-based molecular subtyping of cancer [3,31]. In their approach, expression of a set of 494 breast tumor-cell–intrinsic genes was defined to overcome intra-tumoral cellular heterogeneity, arising primarily from the stroma [3]. Using this strategy, we used previously deposited prostate-tissue cell-lineage–specific gene expression profiles assessed by bulk and single-cell RNA sequencing [17,18]. We discovered four transcriptomic subtypes of primary prostate adenocarcinoma, characterized by prostate cancer-relevant genomic alterations (i.e., SPOP mutation, ETS-family fusions, and PTEN and TP53 deletion/mutations). Our classification partly overlapped with earlier findings from whole-transcriptome-based clustering approaches [9]. However, subtype definitions were not absolute, resulting in classification of ~20% of tumors as mixed. Intriguingly, *KLK3* and ACP3 RNA expression levels, encoding PSA and PAP, respectively, showed potential to identify subtypes; this was further supported by serum PSA and PAP levels measured before RP or docetaxel chemotherapy. Identification of cancer molecular subtypes has deepened our understanding of cancer biology and clinical implications, including therapeutic target identification. In breast cancer docetaxel adjuvant chemotherapy was not beneficial in the luminal A population or in patients with ER-positive and HER2-negative cancers [32–34]. We claim that such findings can be translated to PCa. Our analysis suggests that subtype A, with the strongest AR activity and highest serum PSA levels, should undergo treatment with the new AR target agents [35]. Also, data from recent molecular profiling of mCSPCs support that the SPOP-mutated tumors are less likely to become castration-resistant [36].

The established PCa serum biomarker combination of PSA and PAP may be useful to predict the transcriptomic subtype and docetaxel sensitivity at an advanced stage. Indeed, reports shows that the 5-year survival rate was significantly lower in metastatic cancer patients with a low PSA/PAP ratio than with a high ratio (24% vs. 48%, P=0.002) [37]. In localized tumors, elevated PAP before treatment has been regularly identified as a significant prognostic factor following definitive therapies [38–41]. Although the benefit of docetaxel is repeatedly seen in mCRPC, adjunct docetaxel therapy is not superior to androgen deprivation therapy alone in high-risk cancer with rising PSA only [42]. We postulate that patients with rising PSA only are most likely of subtype A, which is predicted to be insensitive to docetaxel but sensitive to AR inhibition. In other words, patient groups with rising PSA and PAP may be suitable candidates to test the benefit of early docetaxel treatment.

Our finding does not contradict earlier reports that ERG induces taxane resistance in CRPC [43]. Rather, it underscores the importance of radiographic and clinical responses over PSA response in mCRPC cases, where increasing numbers of PSA-low neuroendocrine–like cancers are seen. We argue that ERG-fusion tumors can be subdivided based on PTEN or TP53 loss, and that a combined loss might benefit from a platinum-based regimen. [44]

Interpretation of our data is limited due to the study’s retrospective design and unplanned subset analysis. For docetaxel response analysis, we used pre-chemotherapy serum PSA and PAP levels measured after long-term androgen deprivation, and analysis of the magnitude of the decrease in those markers is not complete. A further prospective trial is warranted, particularly in metastatic castration-sensitive or non-metastatic castration-resistant settings, where the benefit of molecular subtyping and tailored therapies can be maximized.

## ACKNOWLEDGEMENTS

This study was supported by a Korea Health Industry Development Institute (KHIDI) grant for research under the Biomedical Global Talent Nurturing Program of KHIDI (HI19C0723). We thank the patients and pathologists who shared their material and made this study possible. We thank Dr. Byung Ha Chung and Dr. Woo Jin Ko for their support and guidance.

---

**Figure S1.**
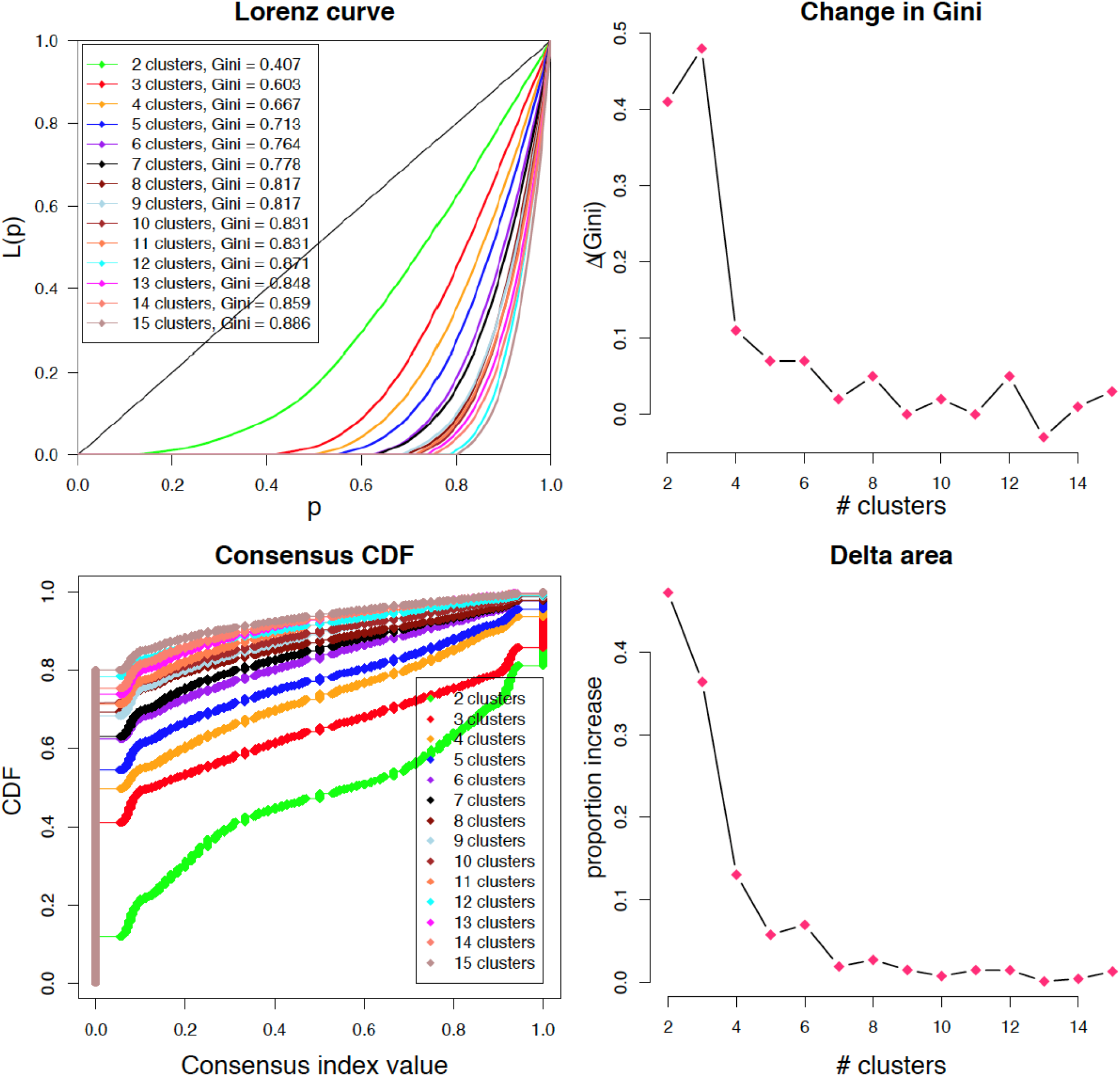
Consensus Clustering of the TCGA-PRAD RNA-Seq Data, Unfiltered.

**Figure S2.**
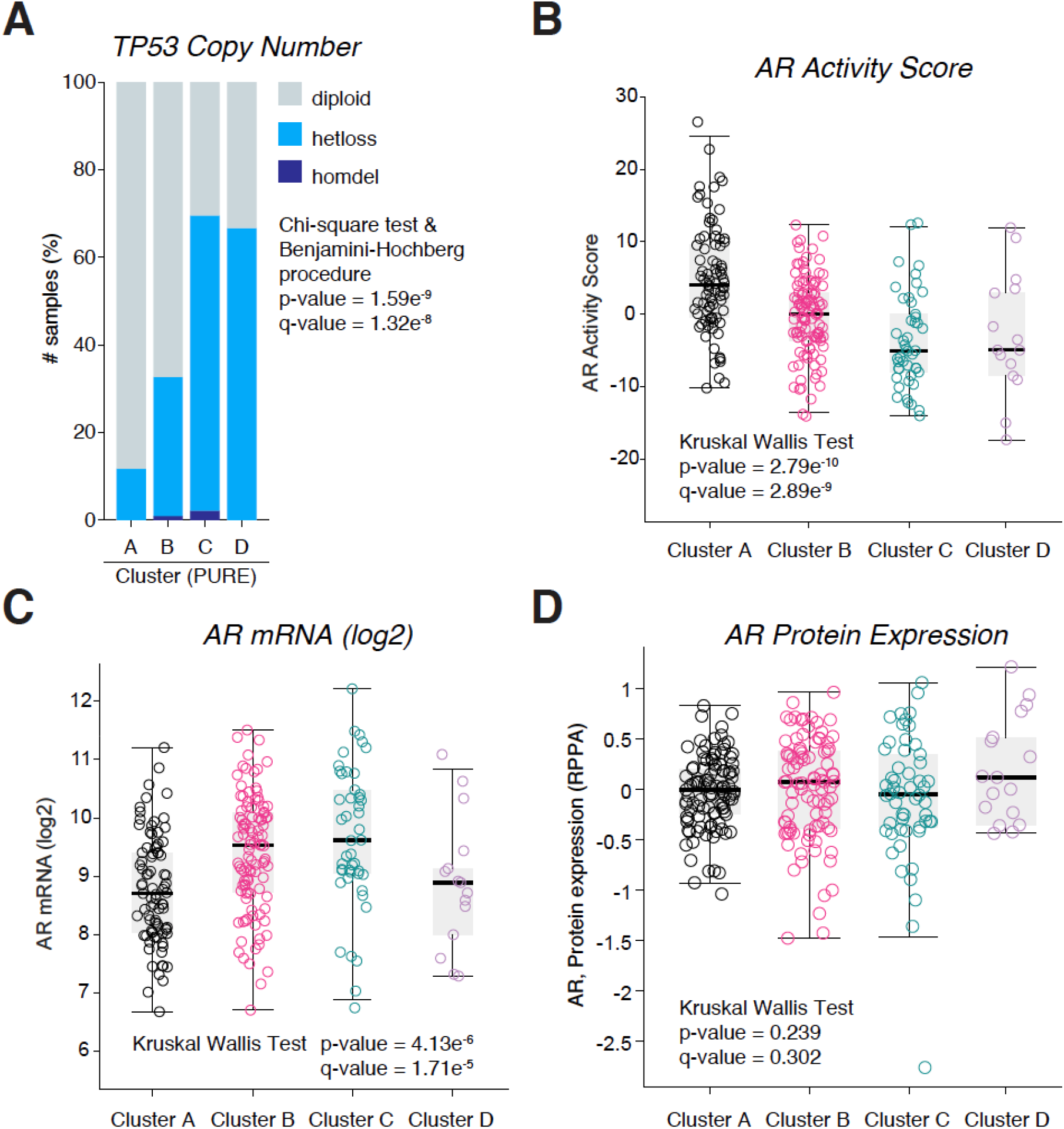
TP53 Copy Number and Androgen Receptor Activity, mRNA and Protein levels of the Four Clusters from the TCGA-PRAD Data. **A.** Status of TP53 copy number of the four clusters, hetloss = loss of heterozygosity;homdel = homodeletion. P and Q values calculated by chi-square test & Benjamini-Hochberg procedure. **B-D.** Androgen receotor (AR) activity score (B), mRNA level (C) and protetin level (D) comparison among the four clusters. P and Q values calculated by Kruskal Wallis Test. All original data downloard from the cBioPortal.org.

**Figure S3.**
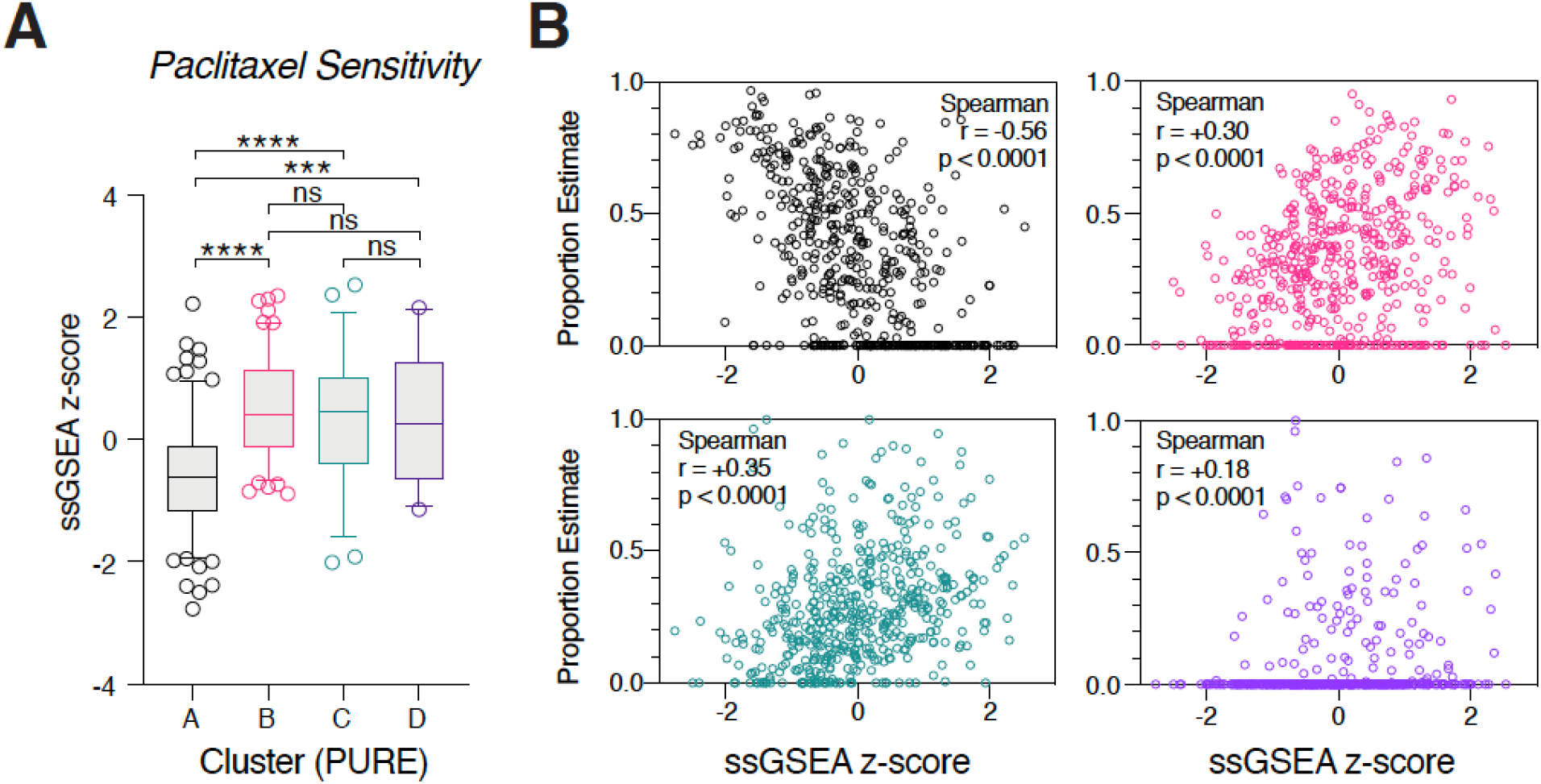
*In Silico* Paclitaxel Sensitivity Test. Scatter plots of the four cluster proportion estimates (PEs, Y-axis) and docetaxel response score of each sample. Blue dot = Cluster A; Red dot = Cluster B; Greed dot = Cluster C; Purple dot = Cluster D. Spearman correlation coefficient and p values are shown in the box.

## Notes

### Competing Interest Statement

The authors have declared no competing interest.

